# Contrastive Learning for Graph-Based Biological Interaction Discovery: Insights from Oncologic Pathways

**DOI:** 10.1101/2024.07.23.604746

**Authors:** Phuong-Nam Nguyen

## Abstract

**Background:** Contrastive learning has emerged as a pivotal technique in representation learning, particularly for self-supervised and unsupervised tasks. Link prediction, crucial for network analysis, forecasts the formation of connections between nodes. Machine learning enhances link prediction by learning patterns from data, leading to improved performance and scalability.

**Method:** In this study, we propose a contrastive learning approach tailored for isomorphic graphs to uncover intrinsic interactions within biological networks. By creating data augmentations through vertex permutations, we train models to learn permutation-invariant representations.

**Results:** In this study, we propose a contrastive learning approach tailored for isomorphic graphs to uncover intrinsic interactions within biological networks. By creating data augmentations through vertex permutations, we train models to learn permutation-invariant representations. Our approach was validated using five cancer-targeting biomarkers: *ADGRF5, TP53, BRAF, KRAS*, and *GNAS*.

**Conclusion:** We discovered new connections between G-coupled receptors (*GPR137B, GPR161*, and *GPR27*) and key pathways, interactions between cyclin-dependent kinase inhibitors (*CDKN1A* and *CDK8*) and specific biomarkers, and identified *NFK-BIA* as a central node linking all targeting biomarkers. This study highlights the potential of contrastive learning to reveal novel insights into cancer research and therapeutic targets. The implementation of this project is made available at: https://github.com/namnguyen0510/Contrastive-Learning-for-Graph-Based-Biological-Interaction-Discovery.

## 1 Introduction

Contrastive learning has emerged as a powerful technique in the field of representation learning, especially for self-supervised and unsupervised learning tasks. This method leverages contrasting positive and negative sample pairs to learn discriminative features without requiring explicit labels. Contrastive learning aims to learn an embedding space where similar instances are pulled together and dissimilar instances are pushed apart. The foundational work by Hadsell, Chopra, and LeCun (2006) introduced this concept through the Siamese network, which utilized a contrastive loss function to train a neural network on pairs of similar and dissimilar images to learn useful representations for tasks such as face verification^1^. This approach is an emergent training technique in Computer Vision literature^2–8^.

Link prediction is a crucial task in network analysis, aiming to predict the existence or emergence of links between nodes in a network. Applying machine learning techniques to link prediction has led to considerable improvements in performance. These methods can automatically learn and generalize patterns from data, providing a more robust and scalable solution. We categorize ML-enabled link prediction into two subgroups: (1) feature-based methods and (2) matrix factorization techniques. Feature-based methods involve constructing a feature vector for each pair of nodes and using these vectors as input to a machine learning model^9,10^. Matrix factorization techniques have been widely used for link prediction, particularly in recommendation systems. Matrix factorization decomposes the adjacency matrix of a network into latent factors, capturing the underlying structure^11^. Link prediction in networks, especially using machine learning techniques, is highly valuable for predicting new interactions among biological networks. Biological networks, such as protein-protein interaction networks, gene regulatory networks, and metabolic networks, consist of nodes representing biological entities (e.g., proteins, genes) and edges representing interactions between them.

Contrastive learning has become a prominent representation learning method, especially in graphs and networks. Contrastive learning frameworks have significantly improved learning robust and meaningful node, edge, and graph representations by leveraging the principles of contrasting positive and negative samples. Deep Graph Infomax (DGI), introduced by^12^, is one of the pioneering works in contrastive learning for graphs. DGI maximizes the mutual information between patch representations (local) and summary representations (global) of graphs. It uses a corruption function to generate negative samples by shuffling node features and a discriminator to differentiate between positive and negative pairs. This approach has shown significant improvements in unsupervised graph representation learning tasks. GraphCL, proposed in^13^, applies data augmentation techniques to generate different views of the same graph. These views serve as positive samples, while other graphs in the batch act as negative samples. The method employs a contrastive loss to maximize the agreement between the augmented views of the same graph. CMRL is introduced in^14^, which leverages multiple graph views, such as node attributes and structural information, to learn node embeddings. By contrasting these different views, CMRL captures complementary information and enhances the quality of learned representations. Despite its usefulness, there still needs to be a research gap in contrastive learning for graph-based datasets. A self-supervised framework is proposed in^15^ that integrates contrastive learning with a node classification task. This approach uses contrastive loss to align node embeddings from different augmentations while predicting node labels. Subgraph Contrast introduced^16^, focuses on learning representations by contrasting subgraphs within a larger graph. This method generates subgraphs through random walks and applies a contrastive loss to maximize the similarity between overlapping subgraphs.

This article proposes a contrastive learning approach on isomorphic graphs to uncover the intrinsic interactions within biological networks. We create data augmentation by permuting vertices and train the models to learn permutation-invariant representations of these networks. Our numerical results demonstrate the proof of concept using five cancer-targeting biomarkers: ADGRF5 (GPR116), TP53, BRAF, KRAS, and GNAS. Compared to established targeting pathways, these new biomarkers have yet to be extensively studied in the clinical literature. First, we discovered new connections between G-coupled receptors, including GPR137B, GPR161, and GPR27, with the GNAS and BRAF pathways. Second, we identified interactions between cyclin-dependent kinase inhibitors, such as CDKN1A and CDK8, with GNAS and TP53, respectively. Finally, we found that NFKBIA is involved in a pathway linking all the targeting biomarkers: ADGRF5, TP53, BRAF, KRAS, and GNAS. NFKBIA acts as an inhibitor of NF-*κ*B, which has numerous therapeutic applications.

This article is organized as follows:

1. Section 2 presents the proposed framework.
2. Section 3 reports the experimental design and result analysis.
3. Section 4 further investigates the mutual exclusivity of found and targeting biomarkers studied across ten databases. We also provide a comprehensive trend analysis of clinical interests regarding the new biomarkers.
4. Section 5 concludes the work by proposing several research extensions.

## 2 Method

### 2.1 Overview

In our proposed framework, we diverge from existing works^12–16^, by focusing on the intrinsic representations derived from the isomorphism of the original graph. While a comparable approach can be seen in GraphCL, which emphasizes data augmentation through techniques such as node dropping, edge perturbation, attribute masking, and subgraph extraction, our method solely employs creating different isomorphic graphs via permutation. This approach is motivated by two key factors. First, methods like node dropping, attribute masking, and subgraph extraction may overlook essential markers during learning. At the same time, edge perturbation may introduce incorrect interactions by randomly adding or dropping edges. Second, our approach compels the model to learn permutation-invariant representations, thereby uncovering intrinsic interaction patterns. The details of our proposed framework will be discussed in Section 2.2, with a theoretical interpretation provided in Section 2.3.

### 2.2 The Proposed Framework

**Model Training:** Let **A**_*n×n*_ be the adjacency matrix associated with a weighted biological network ℊ = (*V, E*), where the node set *V* = {*p*_1_, *p*_2_, …, *p*_*n*_} is the set of proteins with |*V*| = *n* and the edge set *E* is weighted by relevant score between protein pairs. Specifically, we have

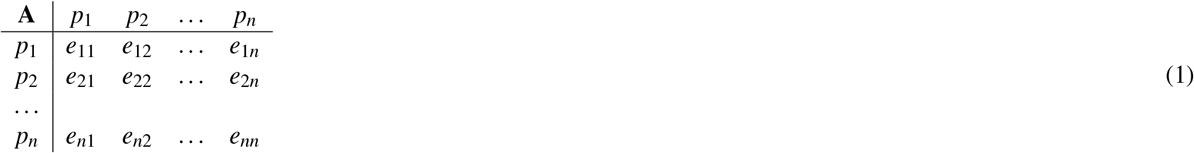

where *e*_*ii*_ = 0 (diagonal elements) and *e*_*ij*_ = *e* _*ji*_, *i≠j* (symmetric graph). An *l*-layer neural network ℱ_Θ_ with model weight Θ = (Θ_2_, Θ_2_, … Θ_*l*_), for which Θ_*k*_ has the structure Θ_*k*_ = (*W*_*k*_, *b*_*k*_) (*k* ∈ {1, 2, … *l*}). For each forward pass, we create two versions of **A** by permutation as data augmentation on the graph, denoted as **A** ↦ (**A**_**1**_, **A**_**2**_). The embedded representation of **A**_*i*_ (*i* = *{*1, 2*}*) is computed as

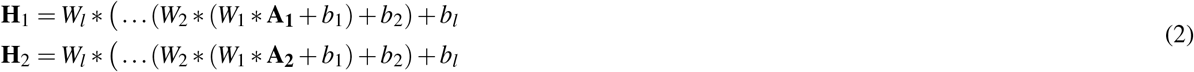

where * is point-wise multiplication. Clearly, we cannot guarantee **H**_*i*_ is a symmetric matrix; thus, we use post-processing the induced representations as follows

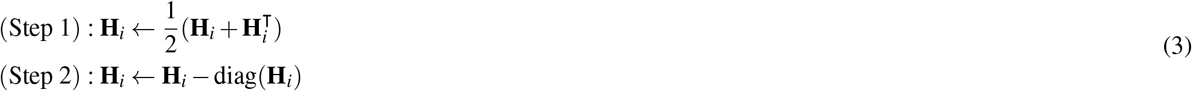

The first step ensures symmetry, and the second removes self-interaction as in the original matrix **A**. We aim to maximize the agreement between **H**_1_ and **H**_2_ as in a conventional contrastive learning framework. We use the objective function ℒ:= (MSE, MAE) and the optimization problem is

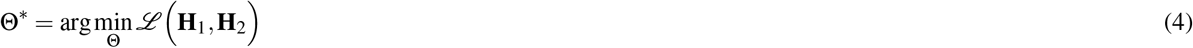

**Model Inference:** We compute **Ĥ** = ℱ_Θ *_ (**A**) to predict the interaction between protein nodes. The Softmax function is applied **Ĥ**←SoftMax(**Ĥ**) to normalized the interaction score to [0, 1] before symmetrizing using Equation 3.

**Remarks:** We use the point-wise multiplication * to avoid saturated embedding **H**_*i*_ and then avoid representation collapsing. Specifically, if the matrix multiplication *W* · **H** is used in Equation 2, this forward pass is attention learning since all other protein interactions are attended in a protein embedding. Consequently, such a message-passing mechanism creates similar embeddings as we increase the number of layers in the model. We want to compute independent latent maps for such interactions and force the model to maximize their agreements. A limitation of our approach is the model complexity, which is *T* (*n*) = *l* ·2*n*^2^ ∈ 𝒪(*n*^2^), where *l* is the number of layers and *n* is the number of input genes.

### 2.3 Theoretical Interpretation

To create an isomorphic graph using a permutation via its adjacency matrix, consider a graph *G* with an adjacency matrix *A*. By applying a permutation matrix *P*, which reorders the vertices of *G*, we obtain an isomorphic graph *G*^*′*^ whose adjacency matrix *A*^*′*^ is given by *A*^*′*^ = *P*^*T*^*AP*. Here, *P*^*T*^ is the transpose of the permutation matrix *P*. This transformation ensures that the structure of *G*^*′*^ is identical to *G*, as the permutation merely reorders vertices without altering the graph’s connectivity. We study the interactions through permutations, which the concept of orbits in permutation groups can elucidate. In this context, an orbit of a vertex under a permutation group action consists of all vertices that can be reached by applying the permutations in the group to the vertex. These orbits reveal the symmetry and equivalence classes of vertices under the permutation, providing insights into the structural properties and the equivalence of interactions within the graph. Permutation-invariant representations on isomorphic graphs are crucial as they reveal intrinsic interactions by focusing on the entire graph’s structure rather than the specific subgraph. By ensuring that the representation remains unchanged under any vertex permutation, we can capture the fundamental properties and relationships within the graph. This invariance is vital for learning from graph data, which requires models to recognize and leverage these intrinsic interactions, leading to more robust and generalizable insights.

## 3 Numerical Results

### 3.1 Case Study

In the numerical demonstration, we aim to discover the pathway between ADGRF5 (GPR116) and iconic drivers of cancers (TP53, BRAF, KRAS, GNAS). Of note, *GPR116* is a potential biomarker for the activation of *CTLA4* discovered in our previous work^17^. *TP53*, often called the “guardian of the genome,” is extensively studied for its role in tumor suppression. Similarly, *BRAF* and *KRAS* are key players in cell signaling pathways frequently mutated in cancers, driving substantial research for targeted therapies. *BRAF* encodes a protein known as B-Raf, part of the MAPK/ERK signaling pathway that regulates cell division, differentiation, and secretion. *KRAS* is a member of the RAS gene family, encoding a protein that also plays a role in the MAPK/ERK signaling pathway. *KRAS* acts as a molecular switch, cycling between active and inactive states to regulate cell proliferation and survival. *GNAS* encodes the G*α*s protein, a subunit of the heterotrimeric G protein complex involved in transmitting signals from cell surface receptors to intracellular effectors, primarily through the activation of adenylyl cyclase and the subsequent production of cAMP. Mutations in *GNAS* can lead to constitutive activation of this signaling pathway, contributing to the development and progression of various tumors. *ADGRF5* (GPR116), also known as Adhesion G Protein-Coupled Receptor F5, is a member of the adhesion G protein-coupled receptor (GPCR) family. GPCRs are a large family of cell surface receptors that play key roles in various physiological processes and are involved in many diseases. *ADGRF5* is predicted to facilitate G protein-coupled receptor activity and is involved in processes like glomerular filtration, pharyngeal arch artery development, and surfactant regulation. It is located in the apical part of the cell and is expressed in the genitourinary system, immune system, liver, lungs, and spinal cord vasculature (Alliance of Genome Resources, April 2022). *ADGRF5* plays a crucial role in glucose homeostasis^18^. This biomarker is involved in the release of somatostatin from pancreatic delta cells, as shown in knock-out mouse models. Whole-body deficiency of *GPR116* results in decreased beta-cell mass, fewer small islets, and reduced pancreatic insulin content. Despite these changes, glucose homeostasis is maintained through compensatory mechanisms that modulate insulin degradation. This highlights *GPR116*’s significant role in regulating glucose levels. *ADGRF5* promotes breast cancer cell migration and metastasis, with its absence impairing cell motility and tumor growth, correlating with increased MMP8 expression and a shift in tumor-associated neutrophils towards an antitumor phenotype^19^. Mechanistically, *ADGRF5* enhances RhoA activation and inhibits ERK1/2 activity, reducing C/EBP*β* activation and leading to heightened MMP8 expression, which attracts antitumor neutrophils and reduces TGF-*β*-enhanced cell motility, making *ADGRF5* a promising therapeutic target for breast cancer. Despite its potential, *ADGRF5* has not been studied widely when compared with well-known biomarkers such as *TP53, BRAF* or *KRAS*, illustrated in Figure 3 and 4.

We use the String database^20^ of protein-protein interaction for this evaluation. In the original graph (**SupMat-S1**), only KRAS and BRAF are connected by a direct pathway, while BRAF connects with TP53 through HSP90AA1. Besides, there is not sufficient evidence to show the connection between GNAS to KRAS, BRAF, and TP53, yet GNAS could be co-activated with these drivers in some certain cancers^21,22^. ADGRF5 and GNAS are connected via RAMP2. The input protein-protein interaction network includes 2,785 proteins and 570,110 edges.

### 3.2 Experiment Designs

We use 2-,4- and 8-layer NNs, trained with three loss functions MSE, MAE, and KL divergence under SGD^23^, Adam^24^ and AdamW^25^ optimizers. We use a flat learning rate of 0.3 for each experiment for the optimizer and train the model for 500 epochs. SGD and KL loss could be more effective (not shown) for our framework, resulting in minimal reduced or intractable loss values. Only 18 models, ignoring the intractable models trained under KL loss, are used for further evaluations. We consider the NNs produced by SGD to be random classifiers. In contrast, models trained with Adam and AdamW can learn to maximize the agreement between the augmented representation, as shown in Figure 2. The optimal neural architectures are found at 4 layers, while the lower depth model (2-layer) has much higher contrastive loss or two deep models (2-layer) do not guarantee the performance gain. To improve the robustness of the proposed pipeline, we use two ensemble training techniques, which are mean average and weighted average. Specifically, we use 18 models with the score computed by

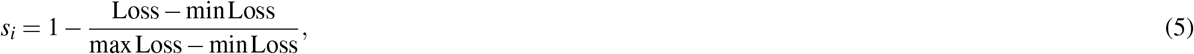

the model with the least loss value will contribute the most to the final synthesized graph. We compute

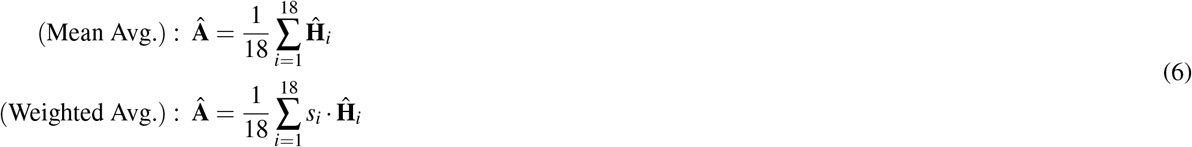

where **Â** is the synthesized adjacency matrix of the predicted protein-protein interaction network.

### 3.3 Evaluation of Synthesized Protein-Protein Interaction

The analysis of the protein interaction networks reveals distinct structural differences between the original, mean-averaged, and weighted-assembled graphs. The original interaction network was filtered to include only significant links within the 98^th^ percentile, resulting in a subgraph of 257 proteins and 275 links. This subgraph differs from the mean-averaged graph, which includes 267 proteins and 280 links, highlighting a slight increase in complexity when interactions are averaged. A noteworthy finding is that 38 protein and 18 links are common between both graphs, indicating a core subset of stable interactions. The graphical illustrations are given in **SupMat-S2**.

Specifically, in the mean-averaged graph, the protein GNAS retains its connection with RAMP2 but loses its direct link with ADGRF5, instead connecting through PDE1A. Three subtypes, PDE1A, PDE1B, and PDE1C, are included in the PDE1 family. PDE1 has potential as a therapeutic target for various diseases, including cardiovascular, pulmonary, metabolic, neurocognitive, renal disorders, cancers, and possibly others^26^. Additionally, GNAS shows a new predicted interaction with GPR161, and KRAS emerges with connections to CTLA4, MRAS, RASA4B, and TP53BP2. TP53 displays novel connections to TP53INP1 and CDK8, with a new indirect link to ADGRF5 via CRBN. CDK8 is a key oncogenic factor, positioning it as a potential tumor biomarker and a promising target for cancer therapy^27^. In the weighted-assembled graph, PDE1A continues to serve as the intermediary between GNAS and ADGRF5, and CRBN remains the bridge for TP53 and ADGRF5. GNAS is also predicted to link to MTOR and CDKN1A, while KRAS exhibits fewer connections, maintaining links only to TP53BP2 and CTLA4. The mTOR pathway is crucial for regulating cell survival, metabolism, growth, and protein synthesis in response to upstream signals in both normal and pathological conditions, particularly in cancer, where it is aberrant signaling, often due to genetic alterations, promotes tumor initiation and progression. Despite extensive use of mTOR inhibitors in therapy, more effective treatments with fewer resistances are needed, with new biomarkers and sequencing technologies enhancing personalized therapy^28^. CDKN1A/p21 is involved in various processes critical for cancer progression, including cell cycle regulation, DNA damage response, apoptosis, senescence, stem cell fate, and EMT. However, its effects can be both pro-tumorigenic and anti-tumorigenic^29^. Both graphs reveal new interactions between BRAF and GPR137B and GPR27, suggesting previously unrecognized pathways. These findings underscore the dynamic nature of protein interaction networks and the importance of different analytical approaches in uncovering novel biological relationships. The GPCR-like lysosomal protein GPR137B controls the localization and activity of Rag and mTORC1^30^. GPR27 is an orphan GPCR that may play a role in neuronal plasticity^31^. We summarize the discovered biomarkers in Table 1.

**Table 1.** Discovered biomarkers by our proposed framework, stratified by targeting biomarkers.

| Category | Genes |
| --- | --- |
| **GNAS-centric** | *GNAS, RAMP2, PDE1A, ADGRF5, GPR161, MTOR, CDKN1A* |
| **KRAS-centric** | *KRAS, CTLA4, MRAS, RASA4B, TP53BP2* |
| **TP53-centric** | *TP53, TP53INP1, CDK8, CRBN, ADGRF5* |
| **BRAF-centric** | *BRAF, GPR137B, GPR27* |
| **Mean average graph** | *NFKBIA, ADGRF5, GNB4, GNAS, ZP2, BRAF, VDAC1, KRAS, BPTF, TP53, RBPJ* |
| **Weighted average graph** | *NFKBIA, ADGRF5, WEE1, TP53, C16orf89, GNAS, NCOA1, BRAF, VDAC1, KRAS, ROCK1* |

### 3.4 Extraction of Significant Sub-graphs for Investigated Pathway

We now extract the most significant sub-graphs that include the targeting biomarkers (ADGRF5, TP53, BRAF, KRAS, GNAS). This is equivalent to applying the longest path problem on both synthesized networks. In the mean average case, we found the pathway: [NFKBIA, ADGRF5, GNB4, GNAS, ZP2, BRAF, VDAC1, KRAS, BPTF, TP53, RBPJ]. In the weighted average case, we found: [NFKBIA, ADGRF5, WEE1, TP53, C16orf89, GNAS, NCOA1, BRAF, VDAC1, KRAS, ROCK1]. The discovered pathways are shown in Figure 1 and summarized in Table 1. Of note, *NFKBIA* pre I*κ*B*α* (nuclear factor of kappa light polypeptide gene enhancer in B-cells inhibitor alpha) is addressed by both assembled methods. I*κ*B*α* can inhibit NF-*κ*B proteins, holding them in an inactive state within the cytoplasm^32^. NF-*κ*B appears to be crucial in regulating the expression of immunoregulatory genes involved in critical illness, inflammatory diseases^33^, apoptosis, and cancer. Specifically, NF-*κ*B is central in controlling the transcription of cytokines, adhesion molecules, and other mediators^34^. Agents inhibiting this pathway, like glucocorticoids and aspirin, can diminish the inflammatory response, while others, such as dominant negative I*κ*B proteins, enhance the effects of chemotherapy and radiation therapy in cancer treatment^35^. A broad range of common human malignancies display abnormal constitutive expression of NF-*κ*B, leading to tumorigenesis and cancer survival in various solid tumors, such as pancreatic, lung, cervical, prostate, breast, and gastric carcinomas^36^. Extensive evidence suggests that NF-*κ*B signaling is primarily involved in the progression of several cancers, which could deepen our understanding of its role in diseases, particularly lung tumorigenesis. Despite the critical role and potential adverse reactions associated with this pathway, various biologics, macromolecules, and small molecules targeting it are available or in trials, with many repurposed drugs also targeting NF-*κ*B, and small molecules comprising 76% of US FDA approvals in the last decade^37^.

**Figure 1.**
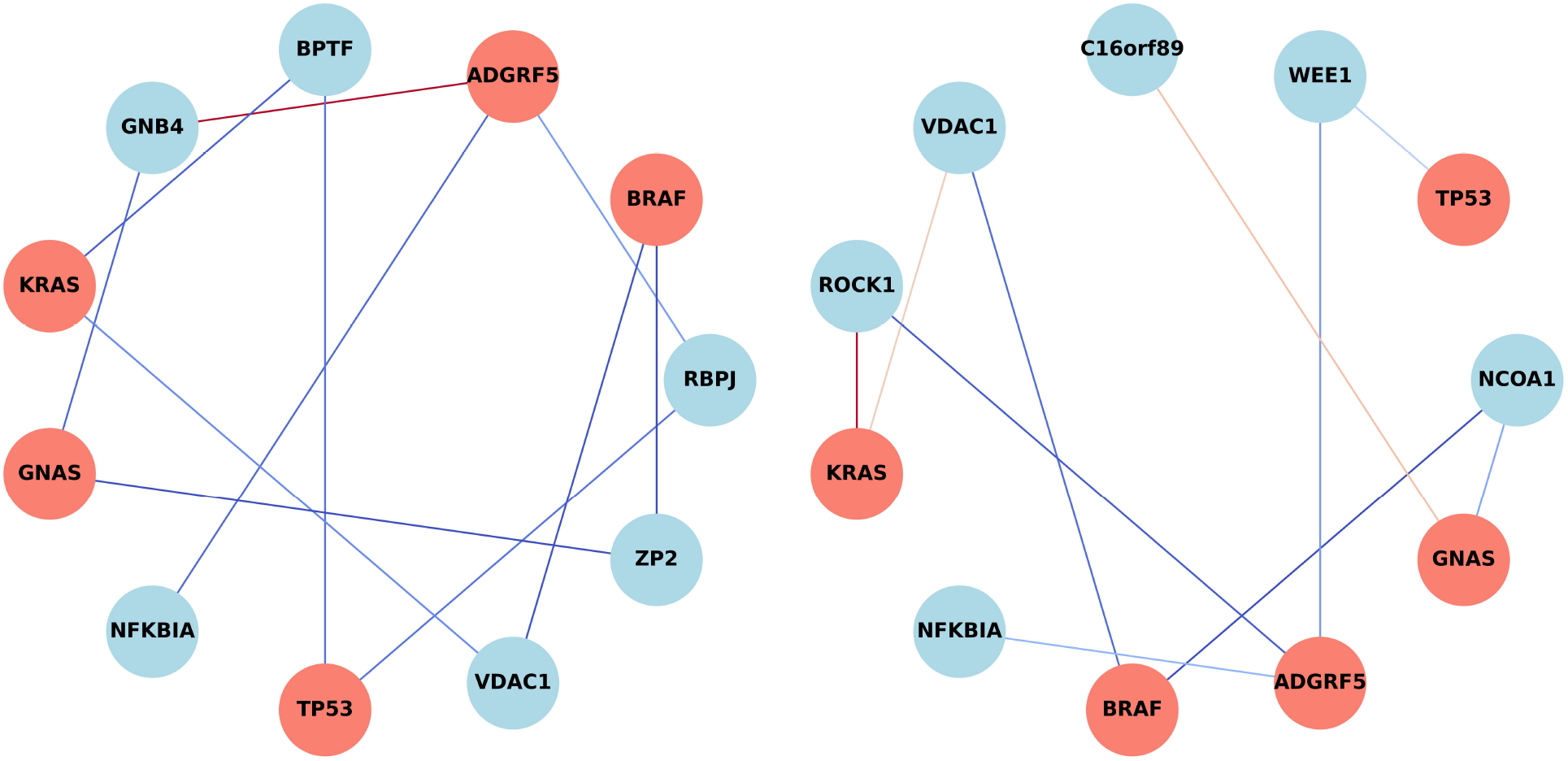
New interaction links discovered by the proposed framework considering targeting pathways (ADGRF5, TP53, BRAF, KRAS, GNAS), left: mean average ensemble, right: weighted ensemble. The score is annotated from low (blue) to high (red).

**Figure 2.**
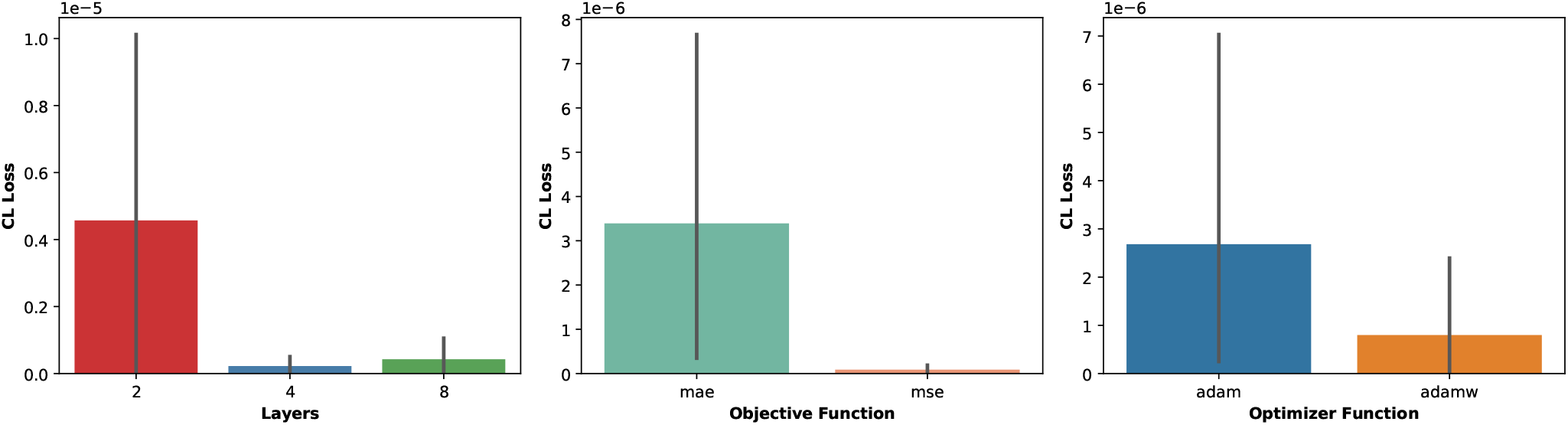
Results of contrastive learning using 2,4 and 8-layer NNs with MSE and MAE loss under Adam and AdamW optimizers

**Figure 3.**
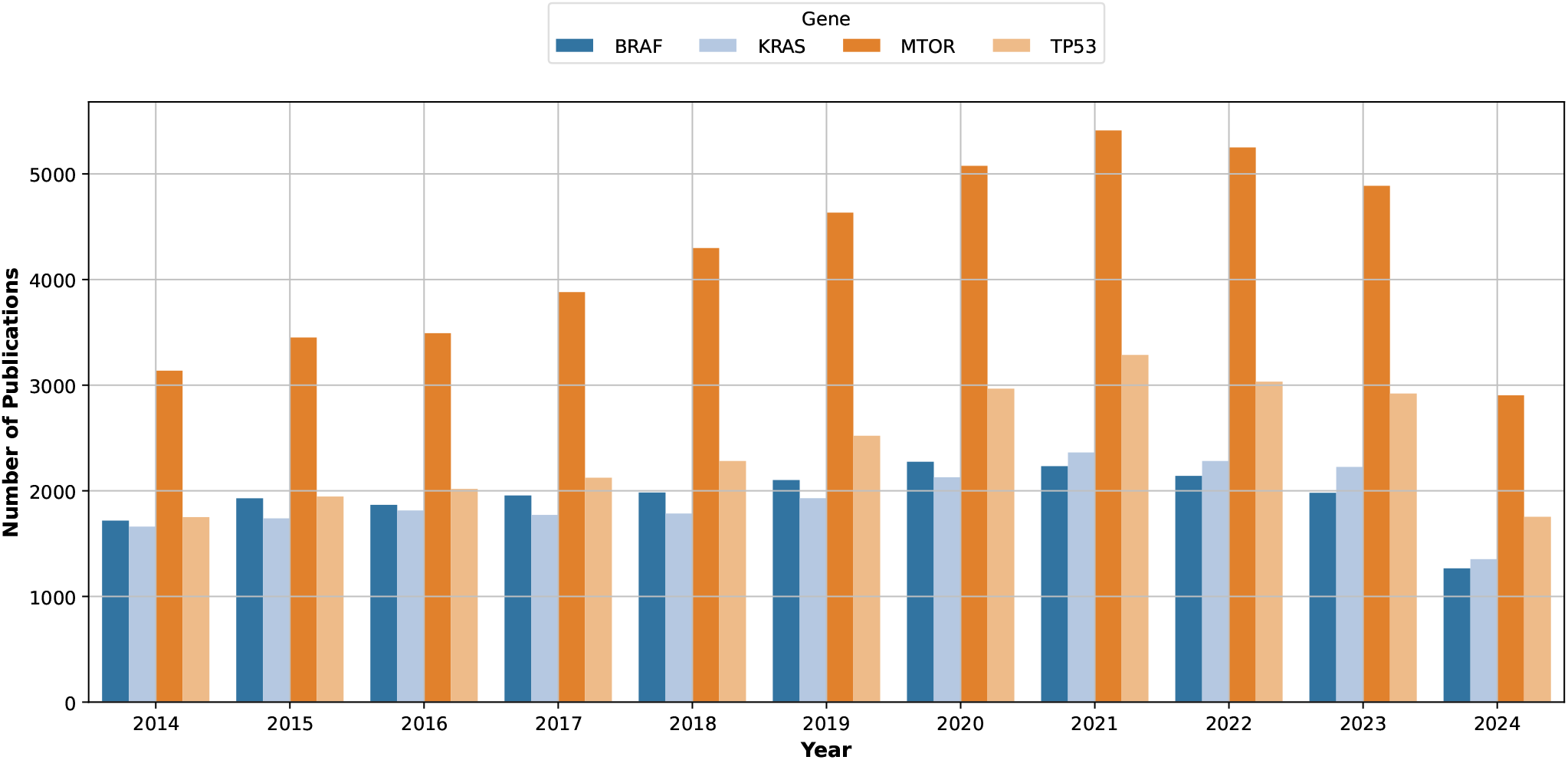
Trends in clinical literature of well-known cancer targets from 2014 − 2024

**Figure 4.**
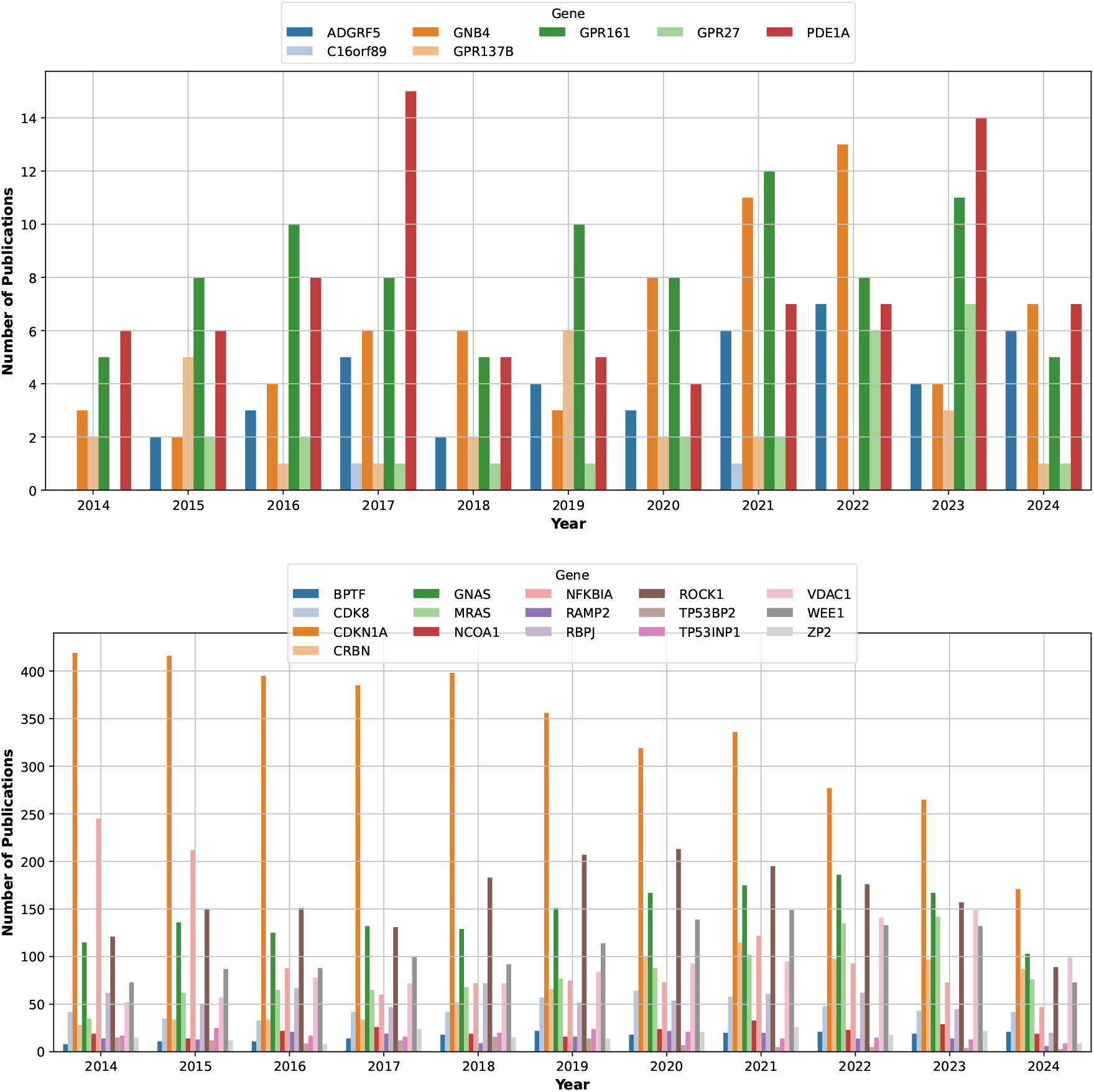
Trends in clinical literature of the discovered targets from. 2014 − 2024. Top: less than 100 papers in total, bottom: more than 100 papers in total.

## 4 Discussion

### 4.1 Mutual Exclusivity Analysis

We investigate the mutual exclusivity of the discovered pathways using the cBioPortal^38^ databases from nine projects^39–47^ with a total of 73,717 samples. We use six queries reported in Table 1. Mutual exclusivity (ME) in cancer genetics refers to the phenomenon where specific genetic mutations rarely or never occur together within the same tumor, suggesting they may perform overlapping roles in cancer development. Co-occurrence in cancer genetics refers to the phenomenon where specific genetic mutations frequently occur together within the same tumor, indicating potential collaborative roles in cancer progression. From our analysis, the ME pairs are (BRAF, KRAS) and (BRAF, TP53). All other pairs are co-occurrences supported by relatively-significant statistical evidence (**SupMat-S3**).

### 4.2 Clinical Interests of The Discovered Biomarkers

Figure 3 displays the number of publications for the genes *BRAF, KRAS, MTOR*, and *TP53* from 2014 to 2024. All four genes show a marked increase in publications over this period, with *TP53* leading significantly in the research volume. This indicates a strong and growing research interest in these genes, critical in cancer biology. *MTOR*, central to cell growth and metabolism, also shows significant publication growth, reflecting its importance in cancer and other metabolic disorders. The upward trend underscores the continued and intensified focus on understanding these genes to develop more effective cancer treatments and personalized medicine approaches.

The top panel of Figure 4 shows the number of publications related to several genes *ADGRF5, C16orf89, GNB4, GPR137B, GPR161, GPR27*, and *PDE1A* in the same decade. The general trend for most genes shows increased publications over the ten years. This suggests growing interest and research in these specific genes, which could be attributed to advancements in genetic research techniques, increasing recognition of their relevance in various conditions, or the rise of personalized medicine approaches. For *ADGRF5*, the trend shows a steady increase in the number of publications starting from around 2015, with a significant spike in the early 2020s. This might indicate discoveries or the emerging importance of this gene in specific medical research areas. For *C16orf89*, it displays a more gradual increase in publications, with a notable rise starting around 2020. This gene’s research may be in its earlier stages of gaining attention compared to others. Publications on *GNB4* have steadily increased, with a more noticeable rise from around 2018 onwards. This could reflect a growing understanding of its role in disease mechanisms or therapeutic potential. Both statistics of *GPR137B* and *GPR161* show a consistent upward trend in publications, indicating ongoing and possibly expanding research interest. Although publications for *GPR27* started lower, a gradual increase suggests emerging research interest. Similar to *GPR27*, there is a steady increase in publications on *PDE1A*, indicating growing research interest over the years. Some of these genes might be implicated in cancer research. For instance, G-protein coupled receptors (GPCRs) like *GPR137B* and *GPR161* are often studied for their roles in cell signaling and cancer.

The bottom panel of Figure 4 illustrates the number of publications related to a set of genes *BPTF, CDK8, CDKN1A, CRBN, GNAS, MRAS, NCOA1, NFKBIA, RAMP2, RBPJ, ROCK1, TP53BP2, TP53INP1, VDAC1, WEE1, ZP2* from the same period. A clear upward trend is observed across most genes, particularly for *CDKN1A, NFKBIA*, and *TP53INP1*, reflecting a substantial increase in research interest. This surge can be attributed to advancements in genomic research and the growing importance of these genes in personalized medicine and cancer genetics. Notably, genes like *TP53INP1* and *CDKN1A*, crucial in cell cycle regulation and apoptosis, show significant increases, indicating their critical role in cancer research and therapy development.

## 5 Conclusion

In conclusion, we introduced a contrastive learning framework on graphs that preserves the intrinsic features by graph isomorphism. Our findings revealed novel connections between G-coupled receptors (GPR137B, GPR161, and GPR27) with the GNAS and BRAF pathways, interactions between cyclin-dependent kinase inhibitors (CDKN1A and CDK8) with GNAS and TP53, and the involvement of NFKBIA in a pathway linking all the targeted biomarkers. Future work should validate these interactions in clinical settings, explore the therapeutic potential of targeting NFKBIA, and enhance the proposed framework to incorporate additional biomarkers and pathways. Furthermore, integrating deep sequencing technologies and developing innovative therapies with better efficacy and less drug resistance is crucial for advancing personalized cancer treatment.

## Supporting information

SupMat-S1

SupMat-S2

SupMat-S3

## Supplemental Materials

- **SupMat-S1:** Original protein-protein interaction networks integrated from String database.
- **SupMat-S2:** Synthesized interaction networks.
- **SupMat-S3:** Mutual exclusivity analysis from ten projects on cBioportal.

## Declarations

### Funding

The research was supported by RF-Bio-N.00001 and RF-AI-N.00007 from G.A.I.A QTech LLC, Vietnam. The author would like to thank Phenikaa university for supports during the project.

### Conflict of interest

The authors declare that they have no conflict of interest.

### Ethics approval and consent to participate

The study was conducted in accordance with ethical standards.

### Consent for publication

All authors consent to the publication of this work.

### Data availability

The data supporting the findings of this study are available upon reasonable request.

### Code availability

The implementation of this project is made available at: https://github.com/namnguyen0510/Contrastive-Learning-for-Graph-Based-Biological-Interaction-Discovery.

## Notes

### Competing Interest Statement

The authors have declared no competing interest.

