## Supplementary figures and images for "Contrastive Learning for Graph-Based Biological Interaction Discovery: Insights from Oncologic Pathways"

### mean_avg_synthesized.jpg

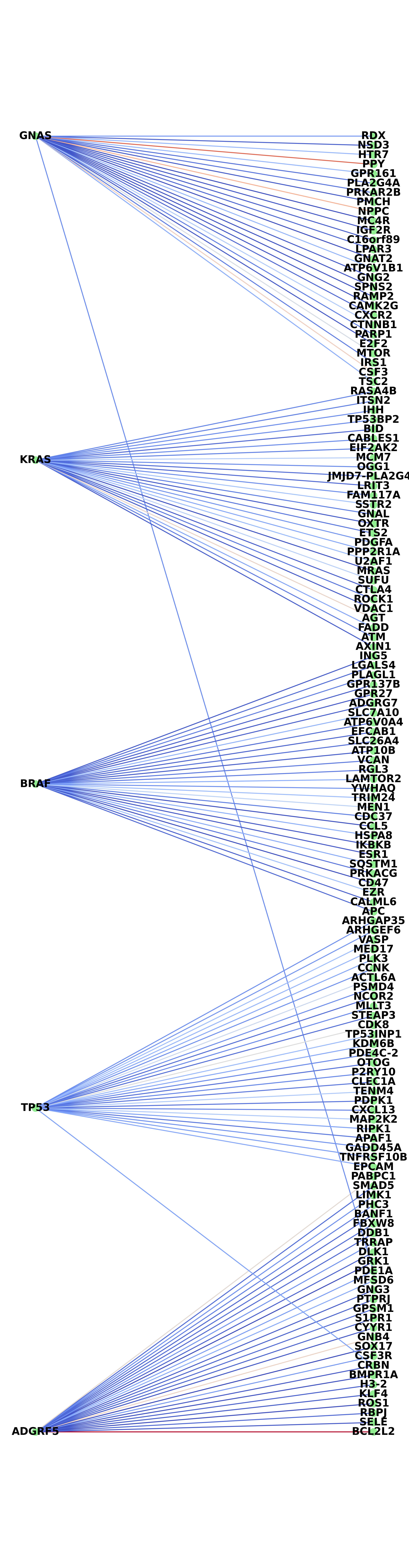

### SupMat-S1_original_graph.jpg

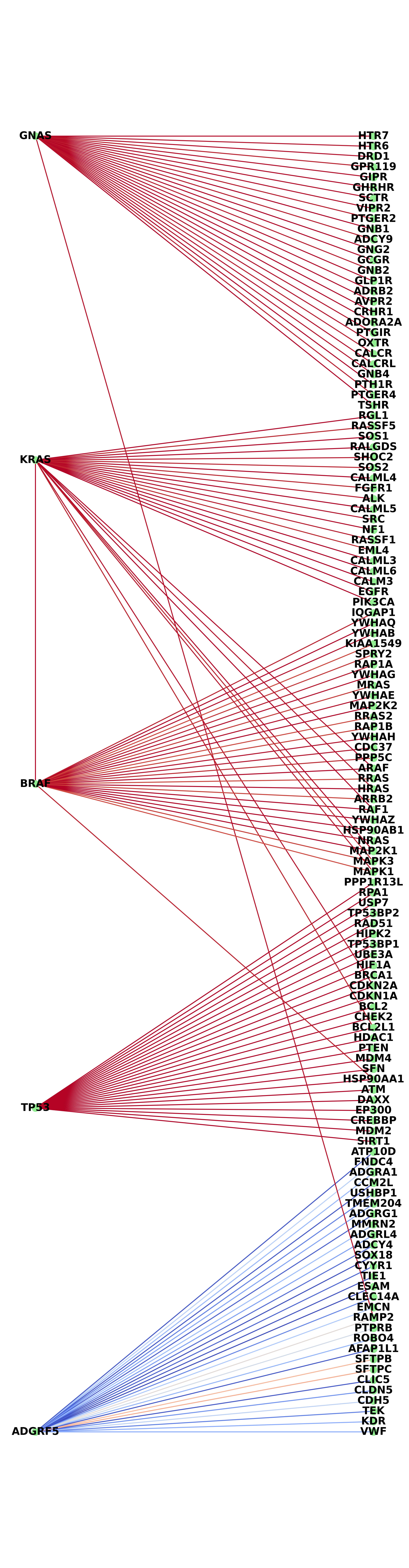

### weighted_avg_synthesized.jpg

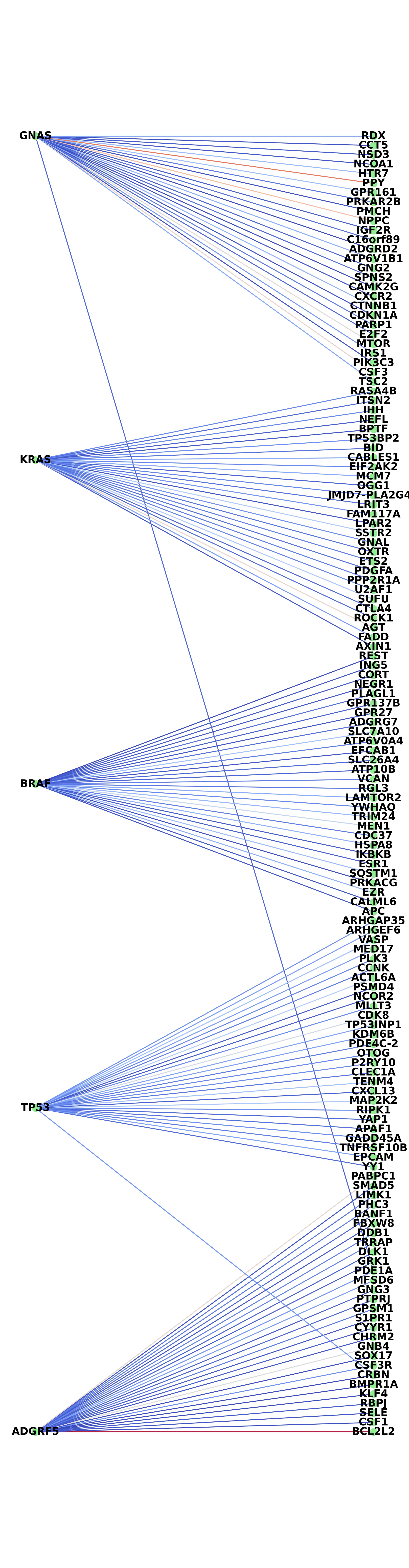
